# Patterns of crossover distribution in *Drosophila mauritiana* necessitate a re-thinking of the centromere effect on crossing over

**DOI:** 10.1101/2024.11.11.623017

**Authors:** R. Scott Hawley, Andrew Price, Hua Li, Madhav Jagannathan, Cynthia Staber, Stacie E. Hughes, Stefanie Williams, Anoja Perera, Rhonda R. Egidy, Amanda Lawlor, Danny E. Miller, Justin P. Blumenstiel

## Abstract

We present a SNP-based crossover map for *Drosophila mauritiana*. Using females derived by crossing two different strains of *D*. *mauritiana*, we analyzed crossing over on all five major chromosome arms. Analysis of 105 male progeny allowed us to identify 327 crossover chromatids bearing single, double, or triple crossover events, representing 398 separate crossover events. We mapped these crossover sites along these five chromosome arms using a genome sequence map that includes the euchromatin-heterochromatin boundary. Confirming previous studies, we show that the overall crossover frequency in *D*. *mauritiana* is higher than is seen in *D*. *melanogaster*. Much of the increase in exchange frequency in *D*. *mauritiana* is due to a greatly diminished centromere effect. Using larval neuroblast metaphases from *D*. *mauritiana* –*D*. *melanogaster* hybrids we show that the lengths of the pericentromeric heterochromatin do not differ substantially between the two species, and thus cannot explain the observed differences in crossover distribution. Using a new and robust maximum likelihood estimation tool for obtaining Weinstein tetrad distributions, we observed an increase in bivalents with two or more crossovers when compared to *D*. *melanogaster*. This increase in crossing over along the arms of *D*. *mauritiana* likely reflects an expansion of the crossover-available euchromatin caused by the reduction in the centromere effect. The pattern of crossing over in *D*. *mauritiana* conflicts with the commonly accepted view of centromeres as polar suppressors of exchange (whose intensity is buffered by sequence non-specific heterochromatin) and demonstrates the importance of expanding such studies into other species of Drosophila.

**Article Summary:** In meiosis, homolog segregation is usually ensured by crossovers. The number and distribution of crossovers is in part regulated by cis-acting factors such as the cis-acting centromere effect, a polar suppression of exchange emanating from the vicinity of the centromere. We use SNP-based crossover mapping to show that in *Drosophila mauritiana*, the centromere effect is greatly reduced on four of the five major chromosome arms. We conclude that the centromere effect differs between *Drosophila mauritiana* and *Drosophila melanogaster* and that the ability to attenuate the centromere effect is not a general property of heterochromatin.

## Introduction

### Genetics functions best as a comparative science – Kenneth W. Cooper

Since its discovery over a century ago in *Drosophila melanogaster*, crossing over has been the subject of extensive study in both meiotic biology and of Drosophila genetics. One of the most active areas of that research focuses on determining the factors and mechanisms that control the number of crossover events and their distribution (Lindsley and Sandler 1977). We seek to understand the *cis* and *trans*-acting elements that control the number and distribution of crossovers, as well as those mechanisms that mediate such processes as crossover assurance (Pazhayam, Turcotte, and Sekelsky 2021) and crossover interference (A. H. Sturtevant 1913, 1915; Berchowitz and Copenhaver 2010). Our goal is to use such knowledge to elucidate the genetic basis for the regional variation in crossover frequencies in the euchromatin (Lindsley and Sandler 1977; Szauter 1984) both within a given genome and between species. Long-standing dogma suggests that such control could be exerted both by the existence of *trans*-acting factors that act on a genome wide basis as well as *cis*-acting processes, such as polar exchange suppression exhibited by centromeres and telomeres.

Several approaches have been employed to understand variation in crossover rates between *Drosophila melanogaster* and closely related species. First, mapping studies in *D. simulans* that used visible markers suggested that the genetic map of *D. simulans* is approximately 30% longer than that of *D. melanogaster* (A. H. Sturtevant 1913, 1915; Barker and Moth 2001). Second, True et al. (True, Mercer, and Laurie 1996) measured crossover rates in *D. mauritiana* by using a large collection of marked transposon insertions in *D. mauritiana*. The position of these insertions was precisely determined by *in situ* hybridization to salivary gland polytene chromosomes to create crossover maps. The crossover frequencies between pairs of these insertions were determined with two-factor crosses and sets of adjacent intervals were summed to determine the distance from any given insertion to the most proximal insertion. For *D. melanogaster* and *D. simulans*, the recombination frequence between a given site and the base of the arm was obtained from standard reference maps. This effort allowed comparisons of the maps of these three species. Together, these studies demonstrated that the total map length of *D. mauritiana* was approximately 1.8 times that of *D. melanogaster*, while the total map length of *D. simulans* was estimated to be 1.3 times that of *D. melanogaster*. Surprisingly, the centromere proximal regions of the euchromatin in *D. simulans* and *D. mauritiana* showed greater rates of crossing over when compared to *D. melanogaster*. This difference was not observed in more distal regions. Third, SNP-based maps of crossover distribution in *D. melanogaster* (Miller et al. 2016) and *D. yakuba* (Pettie, Llopart, and Comeron 2022) have allowed more precise measurements of both crossover density along the five autosomal arms and the type of crossover recovered (single, double, or triple crossovers). Presgraves and collaborators have used a variety of tools to decipher the genetic basis for the observed differences in crossover rates in several species within the melanogaster species group (Brand et al. 2018; Brand, Wright, and Presgraves 2019; Cattani, Kingan, and Presgraves 2012). They concluded that interspecies variation reflects the action of both *cis*- and *trans*-acting elements that determine crossover number and position.

#### The centromere effect

The suggestion by True et al. (True, Mercer, and Laurie 1996) and Brand et al. (Brand et al. 2018) that *cis*-acting elements, such as the centromeres, play a major role in creating inter-species variation in crossover number and distribution is intriguing. In *Drosophila melanogaster*, the term ‘centromere effect’ is used to describe a strong polar suppression of exchange generated by centromeres (Fig. 1). Our current understanding of the centromere effect has been recently reviewed by Hartmann et al. (Michaelyn Hartmann, Umbanhowar, and Sekelsky 2019) and by Pazhayam et al. (Pazhayam, Frazier, and Sekelsky 2024; Pazhayam, Turcotte, and Sekelsky 2021), and so we present only a brief overview here. The centromere effect was first observed by Beadle (Beadle 1932) following his analysis of exchange in translocation heterozygotes and then by Sturtevant and Beadle (A H Sturtevant and Beadle 1936) as part of their analysis of crossing over in *X* chromosome inversion homozygotes. Several subsequent studies both by Yamamoto and Miklos (Yamamoto and Miklos 1977; Yamamoto and Miklos 1978) and by Hawley (Hawley 1980) have supported the prevailing view that the centromere effect is generated by the centromeres themselves with its impact on euchromatin attenuated by the expanse of pericentromeric heterochromatin between the centromere and the euchromatin (see Fig. 1A). (Following Pazhayam et al. (Pazhayam, Frazier, and Sekelsky 2024), we should note that the absence of exchange within the satellite-rich heterochromatin itself is an intrinsic feature of heterochromatin (Mehrotra and McKim 2006), but the centromere effect reflects a polar suppression of crossing over in the centromere-proximal euchromatin. It is proposed that the attenuation of recombination by pericentric heterochromatin is not by a special property of heterochromatin but is merely due to the increased physical distance between the centromere and euchromatin where crossing over occurs.)

**Figure 1.**
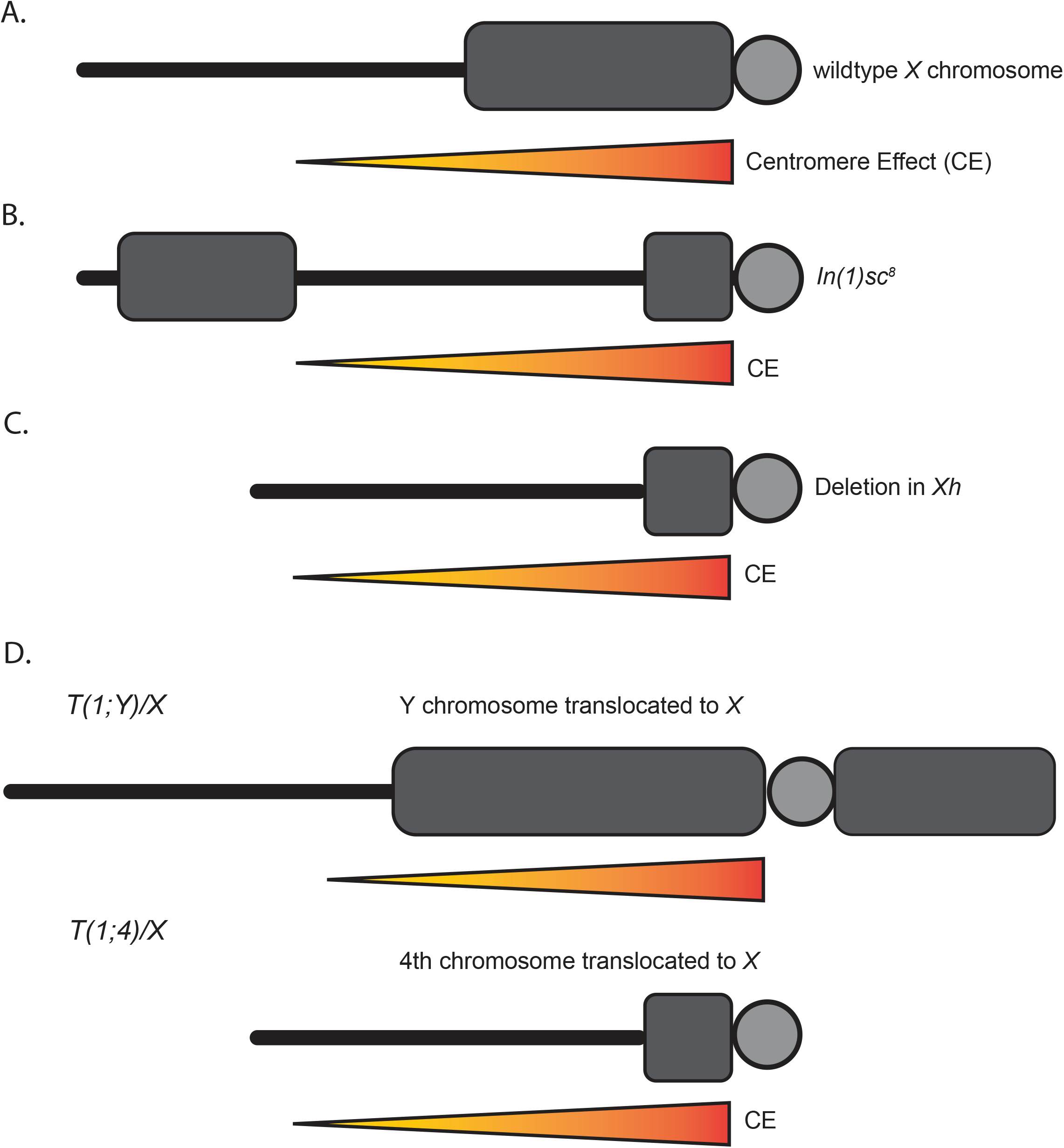
The canonical view of the centromere effect. A schematic depiction of the canonical view of the centromere effect as defined by studies in *D. melanogaster*. Here the centromere effect is depicted as a polar suppression of recombination (orange triangles) emanating out from the centromeres (gray circles). (A) The strength of the centromere effect on exchange in the euchromatin (black lines) is thought to be attenuated by blocks of pericentromeric heterochromatin. (B) As shown by Sturtevant and Beadle (1936), the movement of a large portion of the heterochromatin to a distal site by homozygous inversions allows the centromere effect to penetrate more deeply into the proximal euchromatin. (C) Deleting proximal heterochromatin allows the centromere effect to spread further into the euchromatin (Yamamoto and Miklos 1977; Yamamoto and Miklos 1978; Schalet and LeFevre Jr 1976). (D) *T(1;4)s* show much stronger centromere effects on exchange when heterozygous with a normal sequence *X* chromosome than does *T(1;Y)s* with similar breakpoints on the *X* chromosome. This was presumed to be a consequence of the ability of the larger blocks of heterochromatin on the *Y* chromosome to attenuate the centromere effect more effectively than the much smaller amount of heterochromatin on the *4*^*th*^ chromosome (Hawley 1980).

Two lines of evidence suggest that modifying the extent and severity of the centromere effect might be a strong driver of interspecies variation. First, the work by Brand et al. (Brand et al. 2018; Brand, Wright, and Presgraves 2019) provides evidence that rapidly evolving *trans*-acting factors such as MEI-218 might exert their effects by modifying the severity of the centromere effect. Second, the absence of a strong centromere effect on the *X* chromosome of *D. yakuba* (in the presence of strong suppression seen on autosomes arms) suggests that changes in pericentromeric or centromeric regions themselves may underlie interspecies variation in crossover rate and distribution (Pettie, Llopart, and Comeron 2022).

#### Trans-acting factors that mediate interspecies variation in crossover distribution

Candidates for *trans*-acting factors that mediate interspecies variation in crossover distribution may be the proteins encoded by genes with known mutations that alter the number and distribution of meiotic crossovers in *D. melanogaster* (Baker and Carpenter 1972; Brand et al. 2018; Carpenter and Sandler 1974; Page and Hawley 2005; Pazhayam, Frazier, and Sekelsky 2024). Brand et al. (Brand et al. 2018) showed that replacing the *D. melanogaster mei-218* gene with a transgene copy of its *D. mauritiana* ortholog resulted in an increase in crossing over, most notably in the proximal and distal euchromatin. Indeed, the authors suggest the different *mei-218* genes in *D. melanogaster* and *D. mauritiana* control crossover distribution by “modulating the intensity of centromeric and telomeric suppression of crossing over”. In other words, species level differences arise from both the actions of *cis*-acting mechanisms, such as the centromere and telomere effects, and *trans*-acting factors such as the MEI-218 protein, which is thought to be a pro-crossover factor (Brand, Wright, and Presgraves 2019; M. Hartmann et al. 2019; Kohl, Jones, and Sekelsky 2012).

Here we extend this body of knowledge by providing a SNP-based map of crossing over in *D. mauritiana*. We confirm that the frequency of crossing over is increased in *D. mauritiana* compared to *D. melanogaster*. Our data confirm and extend the suggestion of True et al. (True, Mercer, and Laurie 1996) that much of this increase can be ascribed to a greatly weakened centromere effect on all arms except *2L*. This diminishment of the centromere effect increases the area in which crossovers can occur for four of the five chromosome arms in *D. mauritiana*. The diminishment of crossover suppression in proximal euchromatin, in the absence of obvious changes in the amount of centromeric heterochromatin, requires a reconsideration of our current understanding of the centromere effect.

## Materials and methods

### Fly stocks and husbandry

The following *D. mauritiana* wild-type lines were obtained from the National Drosophila Species Stock Center at Cornell University: Le Reduit, Mauritiana –14021-0241.150 (Mau_LR), Mauritiana 2 –14021-0241.05 (Mau_2), Mauritiana 3 –14021-0241.08 (Mau_3), and Mauritiana genome line –14021-0241.151 (Mau_G). *D. mauritiana* stocks were maintained on a 12 hour light/dark cycle on standard laboratory molasses food at 25°C with ∼40% humidity.

#### Husbandry for the D. mauritiana - Drosophila. melanogaster hybrid

All fly stocks were raised on standard Bloomington medium at 25 °C. *D. melanogaster y w* and *D. mauritiana w*^*1*^ (DSSC#14021-0241.60) were used for the mitotic chromosome spreads.

### DNA preparation and sequencing of parental lines

Genomic DNA from several strains of *D. mauritiana*, including the two strains used for the mapping studies was isolated from 20 females (2 × 10 flies/prep) from each line using the Maxwell 16 Tissue DNA purification kit on the Maxwell 16 instrument (Promega). Samples were eluted in 300 μl water containing RNase A at 20 mg/ml. Like samples were combined, extracted once with phenol/chloroform, and precipitated at -20°C. Samples were resuspended in 22 μl of ultrapure water and quantitated on a Qubit fluorometer (Thermo Fisher Scientific). DNA-seq libraries were generated from 500 ng of DNA. The libraries were made according to the manufacturer’s directions for the Illumina DNA Library Prep, (M) Tagmentation (Illumina, Cat. No. 20060060) and Nextera DNA UD Indexes (set A) (Illumina, Cat. No. 20018704) with 5 cycles of PCR. The resulting short fragment libraries were checked for quality and quantity using the Bioanalyzer (Agilent) and Qubit 2.0 Fluorometer (Thermo Fisher Scientific). The libraries were pooled and sequenced as 100 bp paired reads on a P2 flow cell using the Illumina NextSeq 2000 instrument. Following sequencing, Illumina Primary Analysis version NextSeq2K RTA 3.9.2 and bcl-convert-3.10.5 were run to demultiplex reads for all libraries and generate FASTQ files.

After sequencing, raw fastq files were adapter trimmed using Trim Galore (v0.6.10) and aligned to the *D. mauritiana* reference genome *(D_mauritiana*_ASM438214v1; RefSeq) using bwa mem (v0.7.17). Aligned reads were then deduplicated by processing them through samtools (v1.18) fixmate, sort, and markdup with the -r argument. Variant calling was run on these deduplicated reads using deepvariant (v1.5.0) with the Whole Genome Sequence (WGS) model type argument. SNPs detected by deepvariant were filtered by vcftools (v0.0.16) to only retain biallelic, substitution point mutations with a minimum genotype quality of 20. A list of line-specific parental mutations was generated by comparing the filtered variant calls from the three parental samples and dropping identical shared mutations.

### DNA preparation, sequencing, alignment, and SNP calling of individual flies

Among pairwise comparisons, the Mauritiana 2 (Mau_2) and the Mauritiana genome line (Mau_G) were determined to have the largest number of unshared mutations (i.e. found in Mau_2 but not Mau-G and vice versa) across any pair of parental lines and the broadest distribution across the genome (Supplementary Fig. 1). We used Mau_G and Mau_2 heterozygotes for the recombination studies presented below. Mau_2 virgin females were mated to Mau_G males to produce heterozygous female progeny (denoted as 2-G in the sequence file name, see below). Heterozygous virgin female progeny were then mated individually to Mau_LR, Mau_G, or Mau_2 males to produce male progeny bearing one copy of each maternal, potentially recombinant, chromosome. 108 individual male progeny were obtained from all the crosses: 69 of those progeny were obtained from the cross to Mau_LR males; 25 from the cross to Mau_2 males; and 14 from the cross to Mau_G males. These males were starved for 4-6 hours and frozen for downstream analysis. Sequence was successfully obtained from 105 males.

Genomic DNA was extracted from individual frozen males using the Qiagen DNeasy Blood and Tissue Kit (Qiagen #69506) with the following modifications. Single flies were homogenized using a pestle (Kimble #749520-0090 or similar) with a handheld homogenizer (VWR 47747-370) in 180 μl of ATL buffer, after which 4 μl of RNase A [100mg/ml] was added. All other steps followed the standard manufacturers protocol, except DNA was eluted in 2 × 50 μl of 10 mM Tris (pH 8.0). The DNA eluate was then concentrated to ∼50 μl by centrifugation in Microcon DNA Fast Flow centrifugal filters (Merck Millipore, MRCFOR100) to achieve an average yield of 112 ng/fly. DNA was quantified using a Qubit fluorometer. Individual males were labeled to identify the father, the heterozygous mother, and sibling number. Thus, xG_13.2 was the second male isolated from female 13 mated to a Mau_G father. At least three sibling males were collected from each cross. Numbers may not be sequential because not all preps yielded sufficient DNA for sequencing.

Initially, DNA-Seq libraries were generated for 12 samples from 20 ng of genomic DNA, as assessed using the Qubit 2.0 Fluorometer, according to the manufacturer’s directions with 15 minutes of enzymatic fragmentation at 37°C, targeting 200 – 450 bp, followed by a 10-fold dilution of the universal adaptor and 6 cycles of PCR with the NEBNext Ultra II FS DNA Library Prep Kit for Illumina (NEB, Cat. No. E7805S), and the NEBNext Multiplex Oligos for Illumina (96 Unique Dual Index Primer Pairs) (NEB, Cat. No. E6440S) and purified using the SPRIselect bead-based reagent (Beckman Coulter, Cat. No. B23318). Resulting short fragment libraries were checked for quality and quantity using the Bioanalyzer (Agilent) and Qubit 2.0 Fluorometer. Equal molar libraries were pooled, quantified, and sequenced on a Mid-Output flow cell of an Illumina NextSeq 500 instrument using NextSeq Control Software 4.0.1 with the following read length: 8 bp Index1, 151 bp Read1, 151 Read2, and 8 bp Index2. Following sequencing, Illumina Primary Analysis version NextSeq RTA 2.11.3.0 and bcl-convert-3.10.5 were run to demultiplex reads for all libraries and generate FASTQ files. Subsequently, additional DNA from the same 12 samples was prepared in duplicate using 1/10^th^ reaction volumes, 1:30 fold adapter dilutions, and 8 cycles of PCR with the same reagents and conditions.

The additional 93 samples used in the analysis were prepared using the Seqwell plexWell 96 library prep. DNA-seq libraries were generated from 6.5-16 ng of DNA, as assessed using the Qubit Fluorometer. The libraries were made according to the manufacturer’s directions for the seqWell purePlex DNA Library Prep Kit for Illumina Sequencing Platforms (seqWell, Cat. No. 301067) with kit supplied Indexes (seqWell, Cat. No. 301067) with 8 cycles of PCR. The resulting short fragment libraries were checked for quality and quantity using the Bioanalyzer and Qubit 2.0 Fluorometer. Equal molar libraries were pooled, quantified, and converted to process on the Element Biosciences AVITI with the Element Adept Library Compatibility Workflow, following the Adept Rapid PCR-Free Protocol. The converted pool was sequenced on a 2×150 Cloudbreak High Output flow cell (Cat. No. 860-00003), using AVITI OS 2.3.0 with the following read length: 10 bp Index1, 10 bp Index2, 151 bp Read1, and 151 bp Read2. Following sequencing, Bases2Fastq was run to demultiplex reads for all libraries and generate FASTQ files.

### Detection and validation of crossovers

R (R 4.2.3) was used to process VCF files from individual offspring and identify candidate crossover events using parent-specific variants. The first step filters out the offspring vcf file to only retain variants that match the line-specific parental variant list. Each chromosome was then binned into non-overlapping 50 kb bins, summing the number of variants belonging to each of the three parental lines in each bin. For offspring of the Mau_LR crosses, bins in the bottom 25 percentile of the total number of matched variants were dropped. Crossovers were then detected by finding bins where the dominant (most-frequent) non-LR parental line differed from the previous bin. For offspring of the Mau_G and Mau_2 crosses, bins above the 95th percentile or below the 10th percentile were filtered out. In addition to tracking the number of variants belonging to each parental line for each bin, twelve additional features were added, tracking the number of Mau_2 and Mau_G specific mutations found in the three upstream and downstream bins. Kmeans clustering was applied on those 14 total features (current, upstream, and downstream MauG and Mau2 counts) to cluster each bin into one of two clusters. Crossovers were detected by finding bins where the three previous bins belong to one cluster and the three following bins belong to the other. Since all sequenced offspring were males, the *X* chromosome crossovers were found using the Mau_LR crossover detection method for all samples. All crossovers were validated visually in IGV (Robinson et al. 2011). Only one crossover event (a possible double crossover on 3L) failed to validate in IGV. Crossover events were categorized into classes based on the number of crossovers observed per chromatid: zero or non-crossover (NCOs), single crossover (SCOs), double crossovers (DCOs), triple crossovers (TCOs), and quadruple crossovers (4COs).

### Availability of sequence data

All sequences used in this study are publicly available at https://www.ncbi.nlm.nih.gov/sra/PRJNA1168080. The sample names for sequences posted identify the paternal strain used. For example, sample name 2-Gx2 refers to heterozygous female (Mau_2/Mau_G) crossed to Mau_2 males, sample name 2-GxG refers to heterozygous female (Mau_2/Mau_G) crossed to Mau_G males, and sample name 2-GxLR refers to heterozygous female (Mau_2/Mau_G) crossed to Mau_LR males.

### DNA fluorescence in situ hybridization

For mitotic chromosome spreads, larval 3^rd^ instar brains were squashed according to previously described methods (Larracuente and Ferree 2015). Briefly, tissue was dissected into 0.5% sodium citrate for 5–10 min and fixed in 45% acetic acid/2.2% formaldehyde for 4–5 min. Fixed tissues were firmly squashed with a cover slip and slides were submerged in liquid nitrogen until bubbling ceased. Coverslips were then removed with a razor blade and slides were dehydrated in 100% ethanol for at least 5 min. After drying, hybridization mix (50% formamide, 2× SSC, 10% dextran sulfate, 100 ng of each probe) was applied directly to the slide, samples were heat denatured at 95 °C for 5 min and allowed to hybridize overnight at room temperature. Following hybridization, slides were washed three times for 15 min in 0.2× SSC and mounted with VECTASHIELD with DAPI (Vector Labs). The following probes were used for in situ hybridization: (AAGAG)_6_, (AATAACATAG)_3_ and AGGATTTAGGGAAATTAATTTTTGGATCAATTTTCGCATTTTTTGTAAG (359 bp repeat) and have been previously described (Jagannathan et al. 2017). Fluorescent images were taken using a Leica TCS SP8 confocal microscope with 63× oil-immersion objectives (NA = 1.4). Brightfield images were acquired using a Keyence microscope. Images were processed using Adobe Photoshop software.

### Maximum likelihood estimates of tetrad classes

The frequencies of tetrad classes, denoted as Er, estimate the number of tetrads with r crossovers, where E_0_ corresponds to tetrads with no crossovers, E_1_ to those with one crossover, E_2_ to those with two, and so on). Crossover events can be categorized based on the number of crossovers per chromatid. Let X= (*XX*_*nCO*_, *XX*_*sCO*_, *XX*_*dCO*_, *XX*_*tCO*_, *XX*_*qCO*_) represent the counts for zero crossover (nCO), single crossover (sCO), double crossovers (dCO), triple crossovers (tCO), and quadruple crossovers (qCO), respectively. Zwick et al. (Zwick, Cutler, and Langley 1999) demonstrated that the distribution of crossover counts follow a multi-nominal distribution

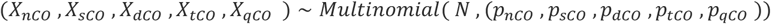

Where *p*_*nco*_, *p*_*sco*_, *p*_*dco*_, *p*_*tco*_, and *p*_*qco*_ denote the probabilities associated with each crossover category, and N represent the total number of observed crossovers. Assuming that crossovers are randomly distributed along chromatids without interference, Weinstein (Weinstein 1936) demonstrated that each crossover class probability is a linear combination of the frequencies of tetrad classes, E_r_. The likelihood function for the observed crossover events is given by:

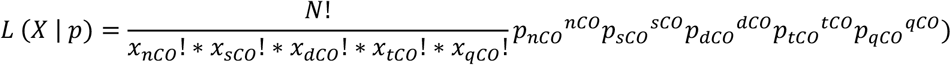

Where

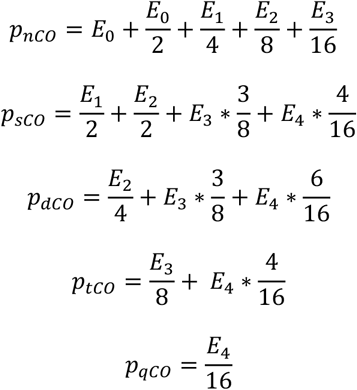

We employed optimization techniques to estimate the maximum likelihood estimates (MLEs) of E_r_s. Specifically, the Nelder-Mead algorithm was utilized via the optim() function in R, with Weinstein’s algebraic solutions serving as the initial parameter values. Additionally, the probabilities were constrained to be positive throughout the optimization process.

A public-facing web app for applying our MLE method on user-submitted crossover count data is available at https://simrcompbio.shinyapps.io/CrossoverMLE/. The app was created using R (v4.3.1) and shiny (v1.7.5) (https://CRAN.R-project.org/package=shiny) and hosted on shinyapps.io.

## Results

### Generation of the Mau_2/Mau_G heterozygous females

In order to choose the two *D. mauritiana* strains used to create the heterozygous females that we used for mapping crossovers, we sequenced four strains of *D. mauritiana* (Mau_G, Mau_2, Mau_3, and Mau_LR) and identified SNPs present in each line (Supplementary Fig. 1). SNPs were filtered using vcftools to only retain biallelic, substitution point mutations with a minimum genotype quality of 20. A list of line-specific parental mutations was generated by comparing the filtered variant calls from the parental samples and removing any identical shared mutations.

We chose to use the Mau_G and Mau_2 lines for our crosses. Mau_G and Mau_2 were chosen because they had the largest number of unshared variants (i.e. found in Mau_2 but not Mau_G and vice versa) across any pair of parental lines. The original analysis found 1,467,376 SNPs unique to one of the two lines, the most out of all possible pairwise comparisons of sequenced strains. The rationale being the more unique SNPs that exist between the parents, the more accurate our ability to position crossover sites. To assist in the identification of the Mau_2/Mau_G recombinant chromosomes in the progeny, the Mau_2/Mau_G heterozygous females were mated in some cases to a third strain, Mau_LR, with a differing SNP pattern.

We then performed single-fly sequencing of 105 individual male progeny from Mau_2/Mau_G heterozygous females. The average depth of sequencing for the initial 12 test samples was 21.42x, while the average depth of the subsequent 93 samples was 15.81x. Analysis of these males allowed us to evaluate crossover events on five chromosome arms (*X, 2L, 2R, 3L*, and *3R*) in each male, obtaining the CO positions for 525 arms. From this data, we identified 198 non-crossover arms (NCOs), and 327 arms with one or more crossover events: 262 single crossover-bearing arms (SCOs), 59 double crossover-arms (DCOs), and 6 triple crossover arms (TCOs). Take together, we mapped 398 crossover events (Table 1) and determined their precise position (Fig. 2). Because our data was relatively low coverage, we did not search for gene conversion events or assay for crossing over on the *4*^*th*^ chromosome.

**Table 1.**
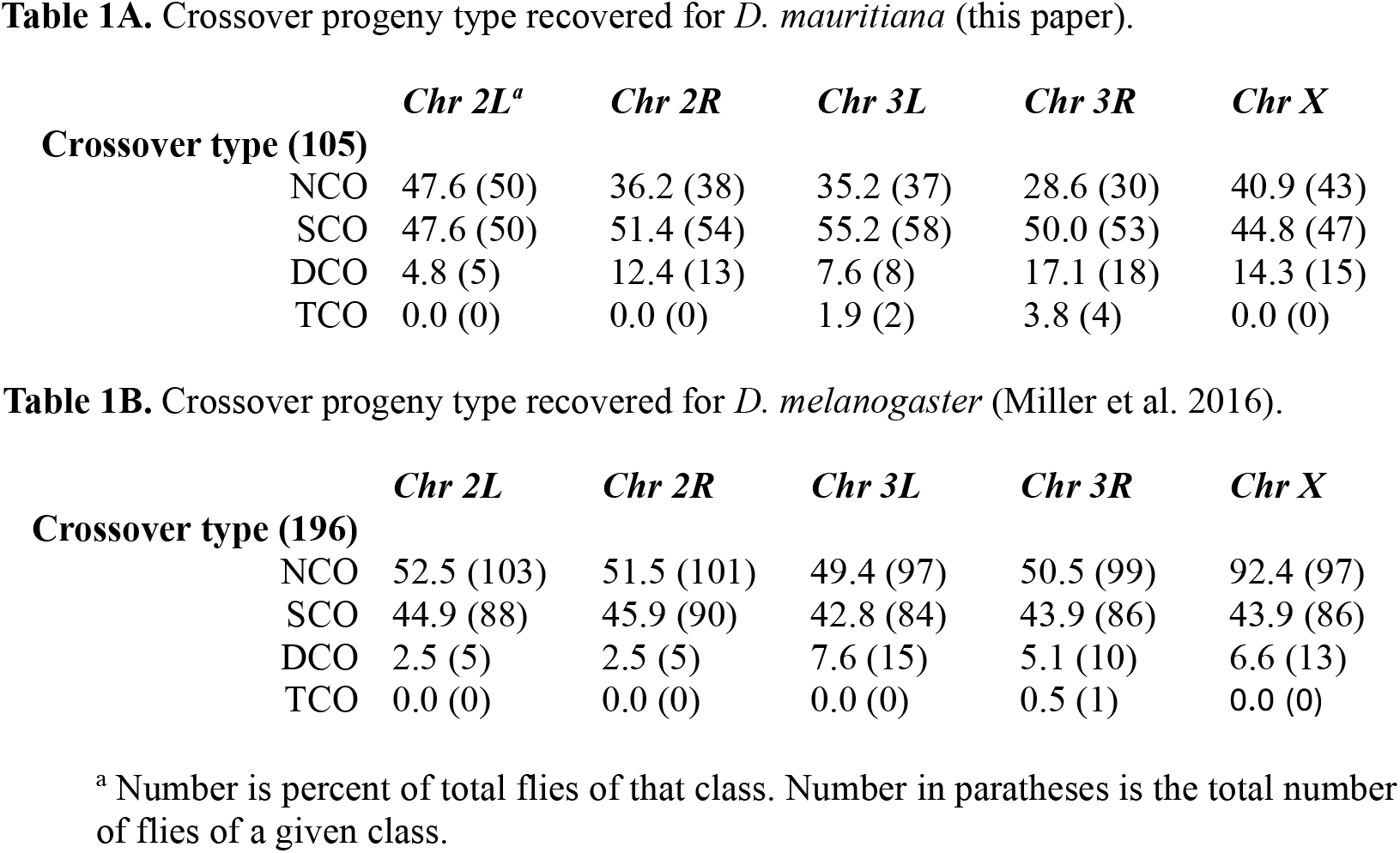
Crossover progeny type recovered.

**Figure 2.**
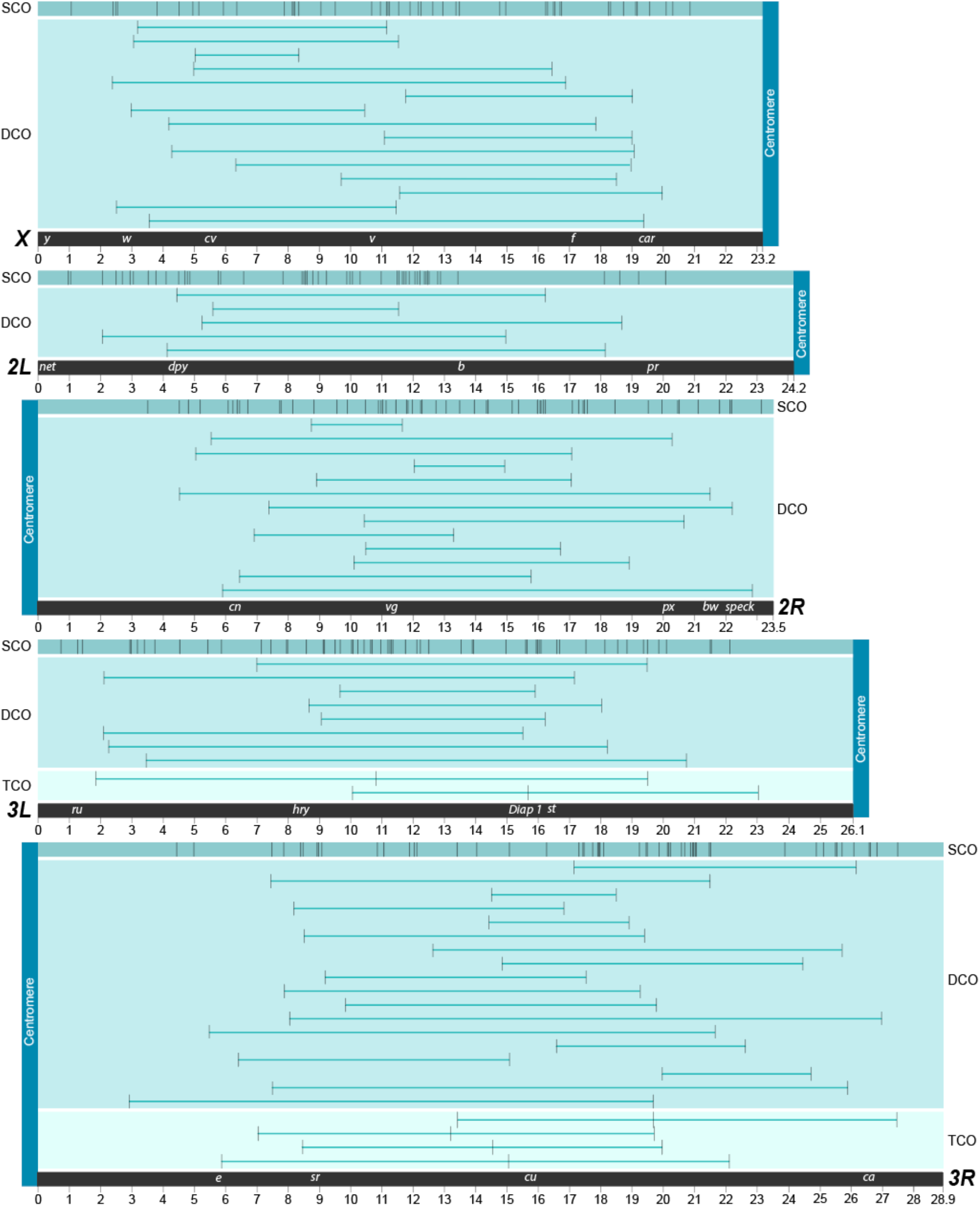
Crossover positions for 398 recombinants. Position of the 398 exchange events observed from 105 individual male progeny, where each panel represents one of the five major *Drosophila mauritiana* chromosome arms (the small *4th* chromosome was not examined). The top track in each panel shows the 262 SCO events. The DCO tracks illustrate the locations and spans of 59 DCOs. Six TCO events were identified, two on chromosome *3L* and four on chromosome *3R*. That the single crossover events making up a DCO or TCO event are well-spaced suggests that interference in *D. mauritiana* is similar to that seen in *D. melanogaster*. Indeed, the average distance between the two crossovers that comprise doubles in *Drosophila mauritiana* is 10.6 Mb, which compares well with the value of 10 MB obtained by Miller et al. (Miller et al. 2016) and suggests that the two species do not differ greatly, if at all, in terms of interference.

### Species-relatedness: a comparison of the genomes

*D. melanogaster* and *D. mauritiana* are both members of the melanogaster species complex. (Supplementary Fig. 2). Their karyotypes and the maps of their salivary gland polytene maps are identical, except for a large paracentric and euchromatic inversion on *3R*. Full genome sequence studies by Chakraborty et al. (Chakraborty et al. 2021) confirm the close relatedness adding only several small inversions on the *X* chromosome to the difference in euchromatic sequence. According to the canonical understanding of the centromere effect, if the pericentromeric regions of heterochromatin in *D. mauritiana* were substantially longer than those of *D. melanogaster*, that should increase centromere-induced suppression of euchromatic crossing over in *D. mauritiana* compared to *D. melanogaster*. However, as shown below, there is no evidence for a substantial difference in the lengths of these regions between the two species.

### Total map length comparison

Table 2 displays the total map length of each arm in *D. mauritiana* compared to the values obtained by Miller et al. for *D. melanogaster* (Miller et al. 2016). In sum, the total map length of *D. mauritiana* is 1.36 times greater than that observed for *D melanogaster*. This increase is substantial, albeit lower than the 1.8-fold increase reported by True et al. (True, Mercer, and Laurie 1996). This increase in map length is not uniform across the five arms. The map lengths of the *3R* and *3L* arms in *D. mauritiana* are increased 1.3 and 1.8 times, respectively, compared to their map lengths in *D. melanogaster*, while the map length of *2L* is increased by only a factor of 1.05. True et al. also found that the greatest increase in total map length was on the *3*^*rd*^ chromosome (2.1 times) compared to *D. melanogaster* than on the *X* (1.8 times) or *2*^*nd*^ chromosome (1.6 times) (True, Mercer, and Laurie 1996).

**Table 2.** Total map lengths (in cM).

| Species | *Chr 2L* | *Chr 2R* | *Chr 3L* | *Chr 3R* | *Chr X* | Total |
| --- | --- | --- | --- | --- | --- | --- |
| *D. mauritiana* | 57.1 | 76.2 | 76.2 | 96.2 | 73.3 | 379.0 |
| *D. melanogaster* | 54.1 | 51.0 | 58.1 | 55.6 | 57.1 | 275.9 |

### Distribution of all crossover events

Fig. 3 presents the crossover density plots for each of the five arms in *D. mauritiana*. Similar plots based on the data from Miller et al. for *D. melanogaster* are presented for comparison (Miller et al. 2016). These plots also show the euchromatin-heterochromatin boundary and the position of a fixed euchromatic inversion on *3R* in *D. mauritiana*. The distribution of medial and distal crossover events in *D. mauritiana* does not appear different from that of *D. melanogaster* (see Fig. 3), even on *3R*. However, there is an obvious increased frequency of exchange in the proximal euchromatin of *D. mauritiana* when compared to *D. melanogaster*. To evaluate whether the increased map length of *D. mauritiana* could be attributed primarily to differences in proximal regions, we performed a statistical comparison with *D. melanogaster* across arm segments divided into thirds: distal, medial and proximal (Table 3). Four (*2R, 3L, 3R* and *X*) out of the five proximal regions showed significantly greater amounts of crossing over in *D. mauritiana*. Only one medial region, on *3L*, showed a significant difference, with *D. mauritiana* also displaying a higher crossover rate. The recombination frequency of two distal regions were significantly different, with one showing a higher recombination rate in *D. mauritiana* (*3R*) and the other showing the opposite pattern (*3L*). Together, these results are most easily interpreted as a decrease in the ability of the centromere effect to reduce the frequency of crossing over in the proximal euchromatin of *D. mauritiana*. This is all the more curious because the expected telomere-adjacent reduction in crossing over is present on all five arms of *D. mauritiana*.

**Table 3.**
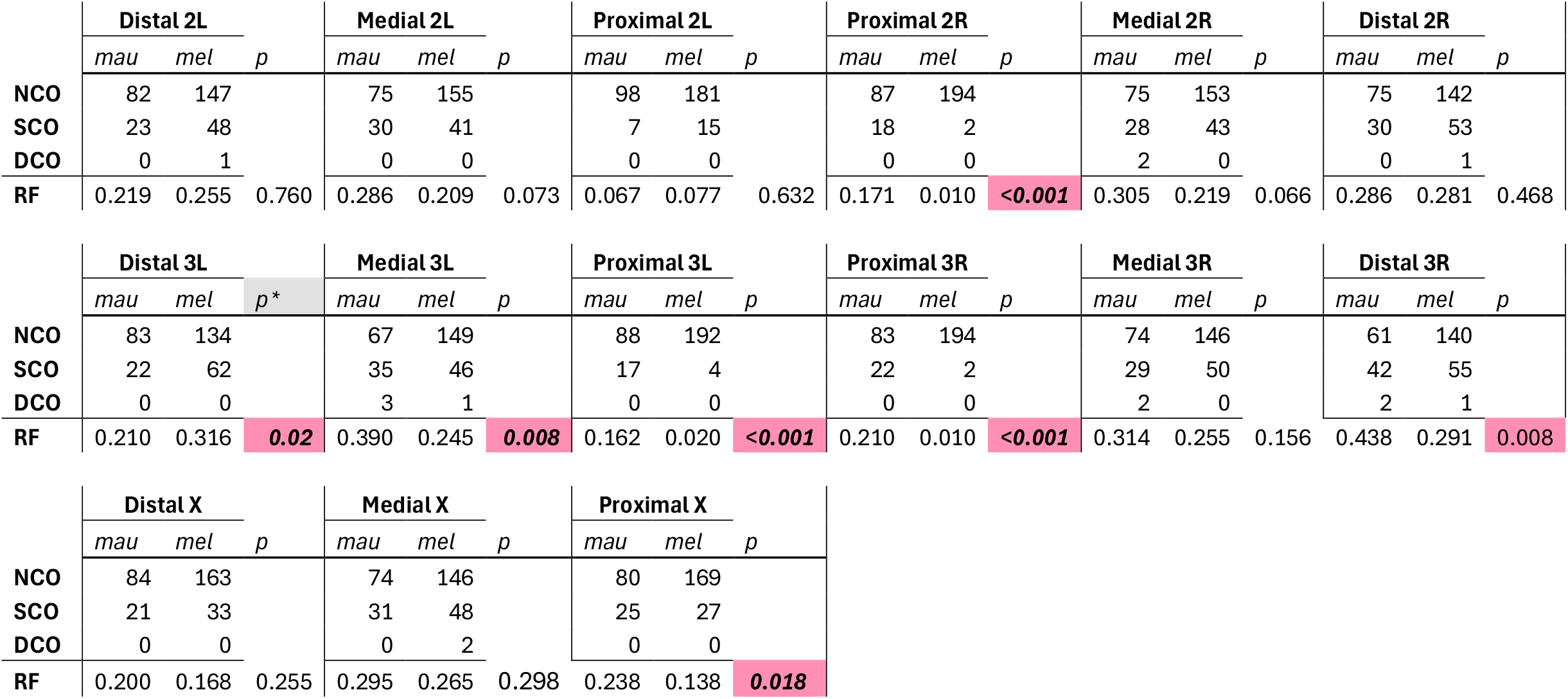
Segment CO status. 1/3 segment (Distal, Medial, Proximal) crossover counts for each chromosome arm (*2L, 2R, 3L, 3R* and *X*). NCO indicates a non-crossover segment. SCO indicates a segment with a single crossover. DCO indicates a segment with a double crossover. RF: Recombination frequency. P-values were calculated by bootstrapping the segment status one million times. P equals the probability, among one million random samples, that the recombination frequency in that segment of *D. melanogaster* is higher than the segment in *D. mauritiana*. *indicates a p-value with the hypothesis that the recombination frequency in that segment of *D. mauritiana* is higher than the segment in *D. melanogaster*. Significant p-values are in pink. Four of five proximal segments show a significant difference, with RFs higher in *D. mauritiana*. One medial segment shows this pattern. Two distal segments show a significant difference between the RF of both species, but in opposing directions.

**Figure 3.**
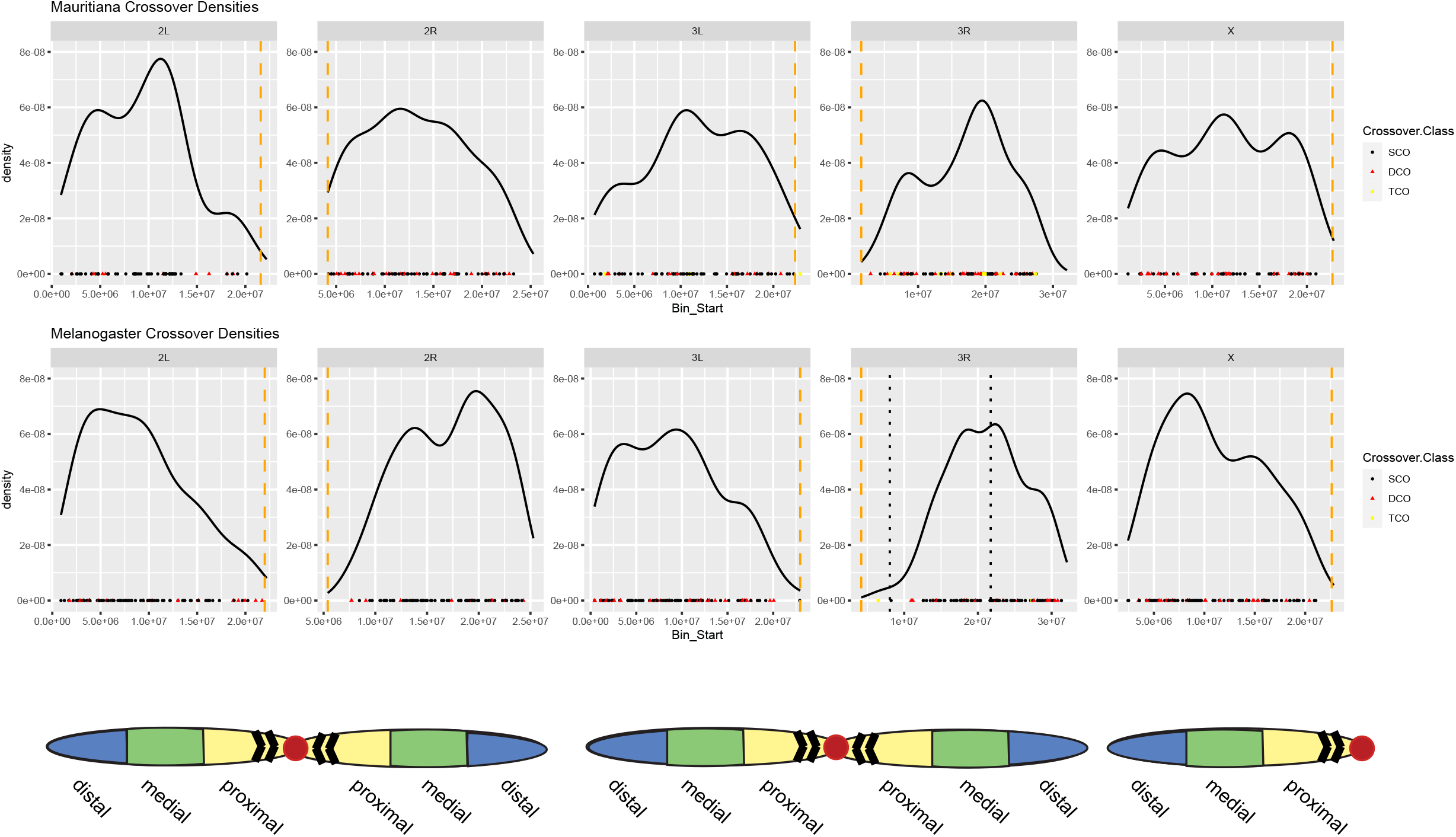
CO density plots for each of the five arms in both *D. mauritiana* and *D. melanogaster*. The euchromatin:heterochromatin boundaries are denoted by dashed yellow lines and the breakpoints of the euchromatic inversion on *3R* are indicated by black dotted lines. The chromosome illustrations indicate the direction of the centromeres in the above plots. Brackets on the chromosomes represent the heterochromatin not included in the plots.

### Dissimilarities between the crossover distributions in D. mauritiana and D. melanogaster cannot be explained by differences in the amount of proximal heterochromatin

One possible explanation for the reduced centromere effect on these four arms might be a substantial increase in the amount of pericentric heterochromatin in *D. mauritiana* that could attenuate the centromere effect relative to *D. melanogaster*. However, careful measures of the genome sizes of these two species shows that *D. mauritiana* is, in fact, slightly smaller than *D. melanogaster* in terms of genome size (Gregory and Johnston 2008), despite the observation by Chakraborty et al. (Chakraborty et al. 2021) that the length of euchromatic sequence is greater in *D. mauritiana* than it is in *D. melanogaster*. A greater amount of euchromatic sequence in *D. mauritiana* in the face of smaller genome size leads to the conclusion that there is *less* heterochromatin compared to *D. melanogaster*.

We also performed DNA fluorescence in situ hybridization (DNA FISH) on mitotic chromosome spreads from *D. melanogaster* and *D. mauritiana* hybrid females. Specifically, we used probes that hybridize with pericentromeric satellite DNA repeats that are abundant in *D. melanogaster* that allow us to identify the species-origin for each mitotic chromosome. Our data clearly show that the *D. melanogaster* chromosomes (arrowheads, Fig. 4 A, B) contain larger DAPI-dense blocks of pericentromeric heterochromatin in comparison to the *D. mauritiana* chromosomes (Fig. 4A, B). These data suggest that alterations in the pericentromeric heterochromatin are unlikely to account for the weakened centromere effect in *D. mauritiana*.

**Figure 4.**
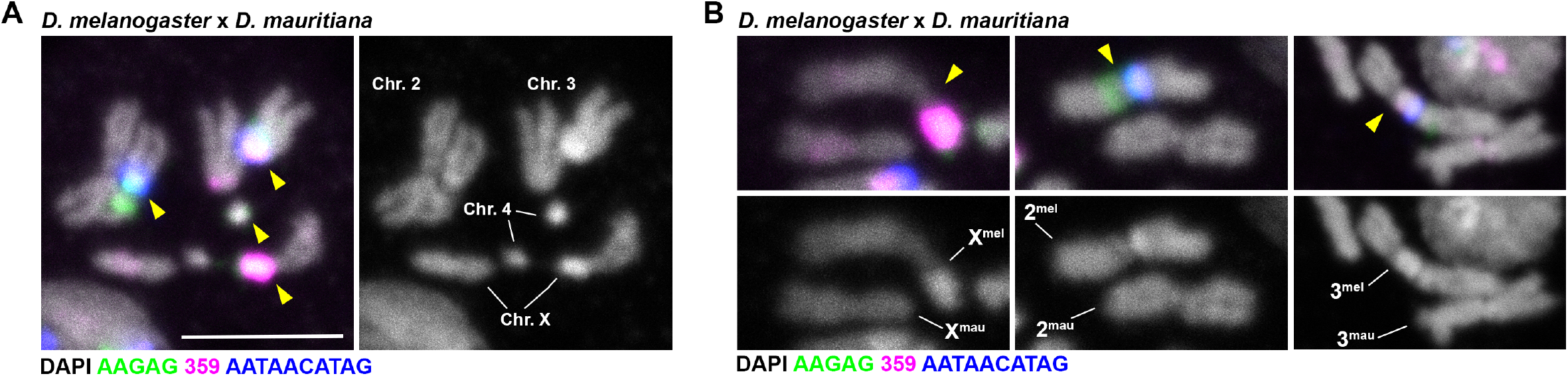
Cytogenetic comparison of the relative lengths of pericentromeric heterochromatin in *D. melanogaster* and *D. mauritiana* hybrids. (A, B) FISH against the (AATAACATAG)_n_ repeat (blue), the (AAGAG)_n_ repeat (green), and the 359 bp repeat (magenta) on larval neuroblast mitotic chromosomes from female *D. melanogaster* and *D. mauritiana* hybrids and costained with DAPI (gray). Arrowheads point to *D. melanogaster* chromosomes.

### A consideration of exchange frequencies at the level of individual bivalents

The crossover density plots shown in Fig. 3 can only be used to identify where the crossovers occurred, not the arrangement of crossovers on individual bivalents. However, we can obtain such information using the data in Table 1A, which presents the primary crossover data in terms of crossover chromatid types (i.e. noncrossover (NCO), single crossover (SCO), double crossover (DCO), and triple crossovers (TCO)). Because each of these types of chromatids can be obtained from bivalents with a higher number of exchanges (Miller et al. 2016), Weinstein (Weinstein 1936) created an algebraic method for characterizing a given population of bivalents in terms of bivalents with no crossovers (E0), one crossover event (E1), two crossover events (E2), three crossover events (E3), and so on. These values are referred to as Exchange Ranks. However, this method can be computationally difficult, especially with small numbers of progeny (Miller et al. 2016; Miller et al. 2018). We used a maximum likelihood estimate to derive Weinsteins values. The method is available online at https://simrcompbio.shinyapps.io/CrossoverMLE/.

Results for this analysis are shown in Table 4. We first consider the *3*^*rd*^ chromosome where in *D. mauritiana, 3R* displays E2 and E3 frequencies of 0.32 and 0.28 respectively, substantially higher than the values of 0.14 and 0.04, respectively, seen in *D. melanogaster*. A similar high frequency of E3 events can be seen for *3L*. Both *2R* and the *X* display an increased frequency of E2 bivalents in *D. mauritiana*. These increases in bivalents with one or more exchange events, came at the expense of non-crossover (E0) bivalents. We admit to being impressed by E0 values of zero for *2R, 3L*, and *3R*. Only on *2L*, the arm on which the centromere effect was unaltered, did *D. mauritiana* show an exchange rank pattern similar to that seen in *D. melanogaster*.

**Table 4.**
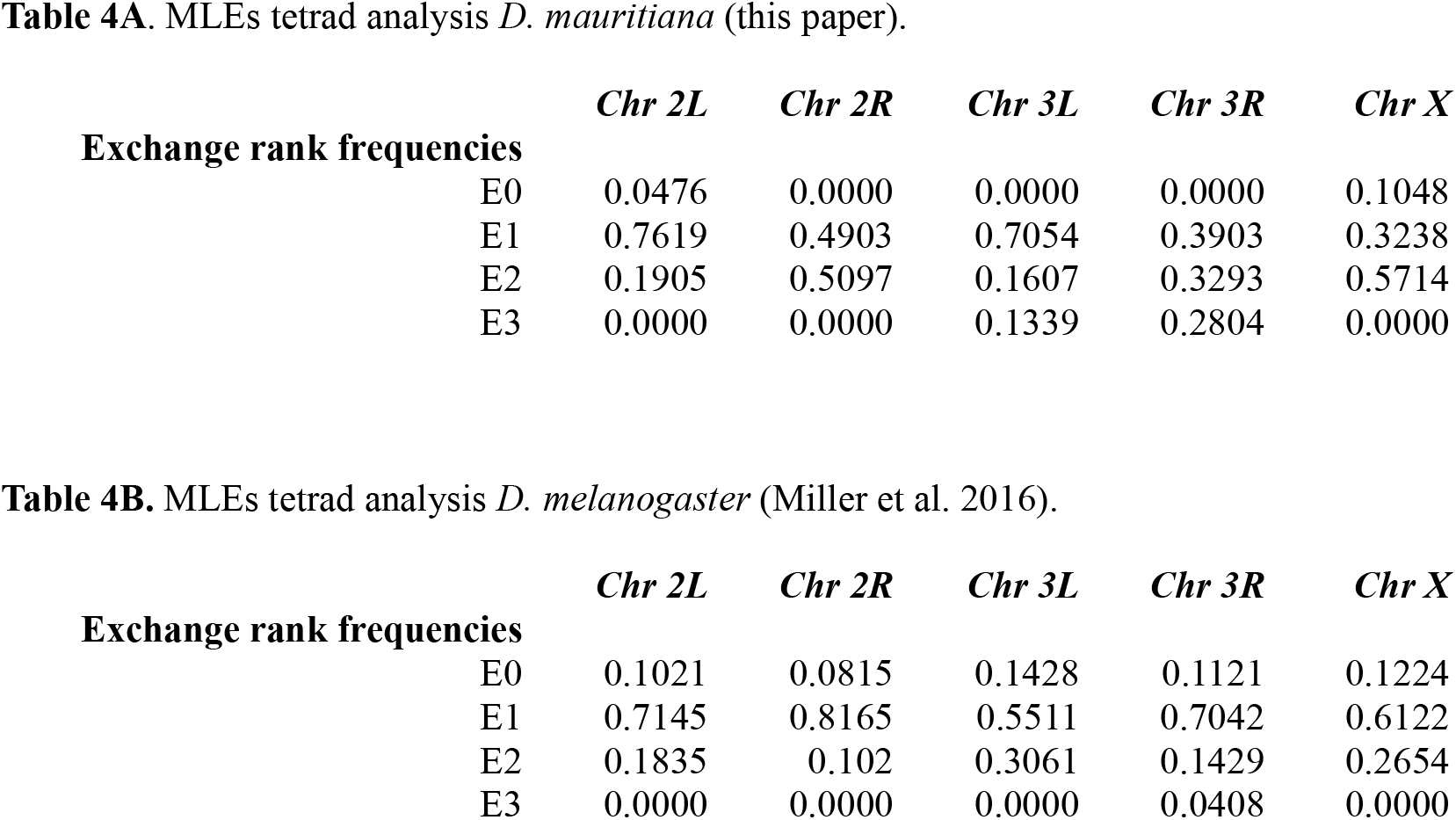
Maximum likelihood estimates (MLEs) of Weinstein tetrad frequencies.

Although more complex interpretations are possible, such as arm-specific alterations in the strength of interference, it seems most parsimonious to explain these data as being primarily driven by a reduction in the centromere effect on each arm. This leads to more of each chromosome arm being used for crossovers, resulting in more double triple and quadruple exchange events. In support of this hypothesis, we note that our average distance between crossovers on DCO-bearing chromatids is 10.6 MB. This compares well with the value of 10 MB obtained by Miller et al. (Miller et al. 2016) and suggests that the two species do not differ greatly, if at all, in terms of interference.

## Discussion

We report both a SNP-based map for crossover distribution in *D. mauritiana* and the development of new and freely available algorithm for determining Weinstein Exchange Ranks using maximum likelihood estimation. At a minimum, our data confirm and extend the work on True at al. (True, Mercer, and Laurie 1996). However, our ability to position each crossover event along a genome sequence denoting the heterochromatin:euchromatin boundaries of each arm allows the additional conclusions regarding the nature of the centromere effect that are presented below. Specifically, we show that despite reduced amounts of pericentromeric heterochromatin on four of the five arms of *D. mauritiana* when compared to melanogaster, the centromere effect on those arms in *D. mauritiana* is much weaker than it is in *D. melanogaster*.

In addition, our ability to detect chromatids bearing double and triple crossovers, and thus determine exchange ranks, allows us to include that the amelioration of the centromere effect on four of the five chromosome arms increase the euchromatin region available for crossing over and thus provide greater opportunity for double and triple crossovers that might otherwise be precluded by interference.

### A new model for the centromere effect

Our data show clearly that reduced heterochromatin on the arms of *D. mauritiana* does not strengthen the centromere effect but, paradoxically, is associated with a decrease. Our data also suggest that the strength of the centromere effect is not related to the simple amount of pericentromeric heterochromatin. Rather, we propose that only some types of heterochromatic regions can buffer the centromere effect, and thus, repeat blocks in the heterochromatin may differ in their ability to attenuate the centromere effect. This suggestion is consistent with the demonstration by Chakraborty et al.(Chakraborty et al. 2021) that one of the most striking differences between the *D. mauritiana* and *D. melanogaster* genomes is the existence of numerous rearrangements within the proximal heterochromatin of the arms.

Finally, Carpenter (Carpenter 1975b, 1975a) has shown that at least in *D. melanogaster*, the synaptonemal complex (SC) appears to have a different morphology, as assessed by serial section electron microscopy, in the heterochromatin than it does in the euchromatin. We wonder if a less obvious perturbation in SC structure might also be found in the proximal euchromatin. It will be interesting to see comparative high-resolution studies of SC structure in multiple species within the melanogaster complex.

### Genes whose protein products may mediate the centromere effect

Our data are interesting in the context of a previous study demonstrating that the replacement of the *D. melanogaster mei-218* gene with its *D. mauritiana* counterpart results in an increase in exchange in the proximal euchromatin in *D. melanogaster* (Brand et al. 2018). (However, as demonstrated by the clear centromere effect observed on *2L* in *D. mauritiana* (see Fig. 3), it cannot be the case that the MEI-218 protein found in *D. mauritiana* cannot impose a centromere effect.) Similarly, *mei-352* mutants in *D. melanogaster* appear to make crossover frequencies proportional to physical distance, thus obviating the centromere effect (Baker and Carpenter 1972; Page and Hawley 2005). We imagine that these proteins, as well as many others, might play roles in controlling the establishment and severity of the centromere effect.

### A centromere effect-based model for the rapid evolution of crossover number and distribution in Drosophila

The rapid change in the structure of the proximal heterochromatin observed by Chakraborty et al. (Chakraborty et al. 2021) may well explain much of the rapid and dramatic change in crossover pattern in the melanogaster complex. Heterochromatic changes can occur without more dire consequences, such as lethality or sterility, while changes in genes that encode trans-acting modifiers of the centromere effect might occur, albeit more slowly, in response to changes in the heterochromatin.

### Concluding thoughts

It is perhaps most surprising to us that *D. mauritiana* does not show a centromere effect on four of the five major arms, but it does show both a level of interference comparable to that observed in *D. melanogaster* and exchange suppression in the telomere adjacent regions. This suggests that changes in the mechanisms that control crossover patterning within a genome (for example, affecting only four of the five chromosome arms) can alter one process, such as the centromere effect, without affecting other processes, such as interference and telomere adjacent suppression. We note in that respect a recent paper showing that the high levels of crossing over and interference observed in budding yeast are not observed in many other species of yeast (Dutta et al. 2024). It occurs to us that a proper understanding of the mechanisms that control crossover placement may well require being willing and able to think outside the well-studied model organism box

## Data availability statement

Original data underlying this manuscript can be accessed from the Stowers Original Data Repository at http://www.stowers.org/research/publications/libpb-2507. All sequences used in this study are publicly available at https://www.ncbi.nlm.nih.gov/sra/PRJNA1168080.

## Acknowledgements

We thank Katherine Billmyre, Monica Colaiacovo, Greg Copenhaver, JJ Emerson, Corbin Jones, Chuck Langley, Nila Pazhayam, Leah Rosin, John True, Kevin Wei, and especially Cathy Lake for helpful discussions and/or comments on the manuscript. We would like to thank Michael Peterson for his work on the sequencing and Mark Miller for his assistance with figure preparation.

## Conflict of interest

D.E.M. is on scientific advisory boards at ONT and Basis Genetics, is engaged in a research agreement with ONT, has received research and travel support from ONT, has received travel support from PacBio, and holds stock options in MyOme and Basis Genetics.

**Supplementary Figure 1. SNP plots for *X, 2L, 2R, 3L*, and *3R* for the four *Drosophila mauritiana* parental strains evaluated for this experiment**. Strains with the highest SNP density compared to the reference were chosen based on analysis of this data. For example, there is a large centromere-proximal region of the *X* chromosome for Mau_3 where SNP density is near 0, which was used to exclude this line from further use. In addition, we compared stocks to maximize inter-strain SNV differences, with Mau_2 and Mau_G showing a high number of unique SNPs between the two stocks.

**Supplementary Figure 2. The melanogaster subgroup**. A dendrogram for the species considered here.

